# Multi-label topic classification for COVID-19 literature annotation using an ensemble model based on PubMedBERT

**DOI:** 10.1101/2021.10.26.465946

**Authors:** Shubo Tian, Jinfeng Zhang

## Abstract

The BioCreative VII Track 5 calls for participants to tackle the multi-label classification task for automated topic annotation of COVID-19 literature. In our participation, we evaluated several deep learning models built on PubMedBERT, a pre-trained language model, with different strategies addressing the challenges of the task. Specifically, multi-instance learning was used to deal with the large variation in the lengths of the articles, and focal loss function was used to address the imbalance in the distribution of different topics. We found that the ensemble model performed the best among all the models we have tested. Test results of our submissions showed that our approach was able to achieve satisfactory performance with an F1 score of 0.9247, which is significantly better than the baseline model (F1 score: 0.8678) and the mean of all the submissions (F1 score: 0.8931).

## I. Introduction

The ever-increasing biomedical literature has posed significant challenges for manual curation and categorization. During the COVID-19 pandemic, manual annotation became even more challenging given the number of COVID-19-related articles growing by about 10,000 per month. This rapid growth has significantly increased the burden of manual curation for LitCovid (1,2), a literature database of more than 100,000 COVID-19-related publications. LitCovid is updated daily with new articles identified from PubMed and organized into curated categories, such as treatment, diagnosis, prevention, transmission, etc. Annotating each article with up to eight possible topics has been a bottleneck in the LitCovid curation pipeline. To support manual curation on topic classification, the track 5 of BioCreative VII calls for a community effort to tackle automated topic annotation for COVID-19 literature.

LitCovid is used by researchers, healthcare professionals, and the public worldwide to keep up with the latest literature of COVID-19 research. Increasing accuracy of automated topic prediction for COVID-19-related literature would be beneficial to both the curators and all the users. The topic annotation in LitCovid is a standard multi-label classification task that assigns one or more labels to each article. The first batch of documents in LitCovid was annotated manually. To support manual curation on topic classification, Chen et al. (2) developed eight deep learning models that integrate embeddings encoded by BioBERT (3) with manually crafted features to predict the probability for topic assignment, one model for each topic. Evaluated on a subset of about 40,000 articles in LitCovid, they were able to achieve an average micro F1 score of 0.81. Jimenez Gutierrez et al. (4) evaluated a number of models on a LitCovid dataset of 8,000 articles and achieved a micro F1 score of around 0.86 with the best performing model using BioBERT.

As a team participating in this task, we evaluated several deep learning models with different architectures based on PubMedBERT (5), a pre-trained language model based on BERT. Our experiments showed that our approach was able to achieve quite satisfactory result (F1 score: 0.9247) that was significantly better than the performance of ML-Net (6), the baseline model using a more general and accessible shallow embedding approach (F1 score: 0.8678). Our method also performed significantly better than the mean F1 score of all the submission (0.8931).

## II. Methods

There are three datasets provided for this task. Based on these datasets, we trained and evaluated several deep learning models with different architectures built on PubMedBERT.

### A. Datasets

The datasets provided for this task included a training dataset of 24,960 articles, a development dataset of 6,239 articles and a test dataset of 2,500 articles. Articles in the datasets contain publicly available metadata of COVID-19-related articles, such as journal names, author names, titles, abstracts, and keywords. Article length vary significantly in terms of the number of sentences in abstracts and the numbers of words in titles and abstracts as shown Figure 1 and Table I.

**Fig. 1.**
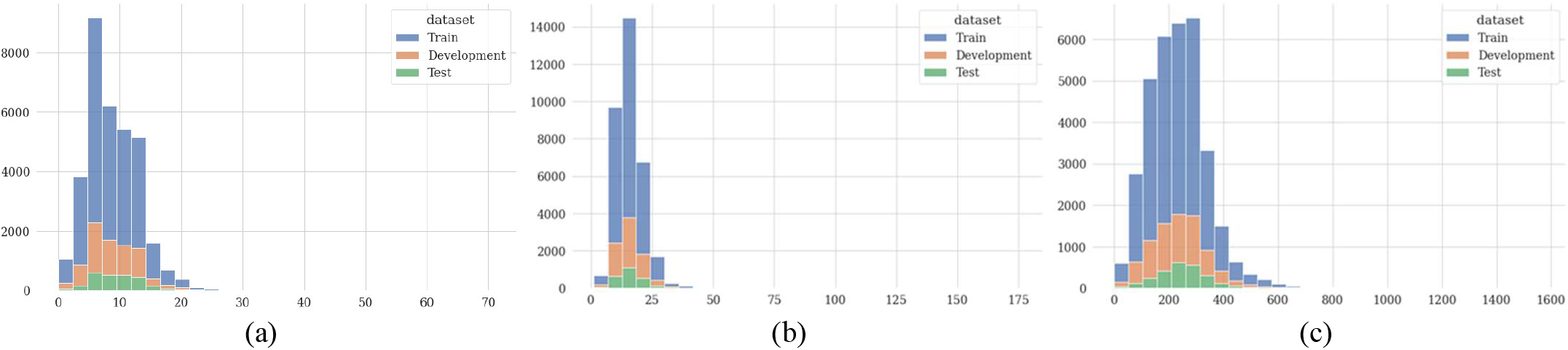
Distributions of (a) number of sentences in the abstracts, (b) number of words in the titles, and (c) number of words in the abstracts in the training, development and test datasets.

**TABLE I.**
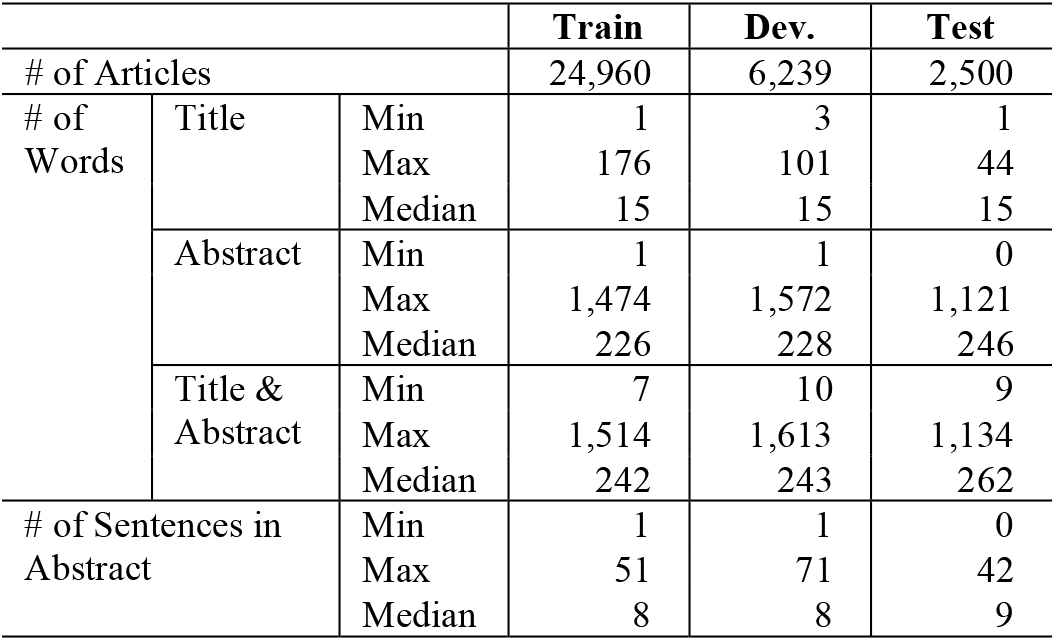
Stateitics of datasets

Each article in the datasets is assigned one or more labels of the 7 topics including Prevention, Treatment, Diagnosis, Mechanism, Case Report, Transmission, and Epidemic Forecasting. Table II shows the breakdown of articles by number of labels and the distribution of each topic in the training and development datasets. While every article can be labelled with multiple topics, more than 95% of the articles in the training and development datasets contain only one (67.4%) or two labels (close to 30%). Distribution of each topic varies significantly from topic to topic. The most frequent topic, namely Prevention, is assigned to more than 40% of the articles and consequently has more balanced positive and negative cases. However, the least frequent topic, i.e., Epidemic Forecasting, is only assigned to around 3% of the articles which results in highly imbalanced positive and negative cases.

**TABLE II.** Labels of the traingin and development datasets

|  | Training |  | Development |  |
| --- | --- | --- | --- | --- |
|  | Articles | Percent | Articles | Percent |
| Total | 24,960 | 100.0% | 6,239 | 100.0% |
| **By number of labels** |  |  |  |  |
| 1 | 16,827 | 67.4% | 4,203 | 67.4% |
| 2 | 7,135 | 28.6% | 1,839 | 29.5% |
| 3 | 860 | 3.4% | 168 | 2.7% |
| 4 | 121 | 0.5% | 24 | 0.4% |
| 5 | 17 | 0.1% | 5 | 0.1% |
| **By different labels** |  |  |  |  |
| Prevention | 11,102 | 44.5% | 2,750 | 44.1% |
| Treatment | 8,717 | 34.9% | 2,207 | 35.4% |
| Diagnosis | 6,193 | 24.8% | 1,546 | 24.8% |
| Mechanism | 4,438 | 17.8% | 1,073 | 17.2% |
| Case Report | 2,063 | 8.3% | 482 | 7.7% |
| Transmission | 1,088 | 4.4% | 256 | 4.1% |
| Epidemic Forecasting | 645 | 2.6% | 192 | 3.1% |

### B. Models

The multi-label classification task can be formulated as: given a document *x* in a collection of ***X***, ***x*** ∈ ***X***, and a finite set of *m* labels ***Y*** = {*y*_1_, *y*_2_,…, *y_j_*,…, *y_m_*}, assign a set of relevant labels ***y*** ⊆ ***Y*** to *x* by learning a classifier *f*: *y* = *f*(*x*).

It is more convenient to identify a set of relevant labels *y* with a binary vector *y* = (*y*_1_, *y*_2_,…, *y_j_*,…, *y_m_*), where *y_j_* = 1 when it is a relevant label and *y_j_* = 0 otherwise. Then to learn the classifier *f*becomes to learn a set of classifiers:

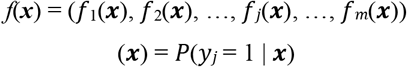

Multi-label classification task is usually considered as a difficult task given the complexity of the task requiring assignment of multiple labels to an article in a large label space. In addition, the large variation in article lengths, the lack of information such as missing title or abstract in some articles, and the imbalanced distributions of topics increase the difficulty of the task.

To tackle the task, we built several deep learning models with different architectures on top of PubMedBERT, a transformer-based pre-trained BERT (7) language model. PubMedBERT was pre-trained from scratch with corpus developed from PubMed articles and it consistently outperformed all the other BERT models in most biomedical natural language processing tasks (5). On top of this model, we applied different strategies to address the aforementioned difficulties. After experimenting with different models trained and validated on the training dataset and evaluated on the development dataset with different strategies, we submitted predictions of five representative models for the test dataset. These five models include:

- The **BERTBASE** model is an architecture with the classifiers built directly on top of PubMedBERT. The title and abstract of each article were concatenated and fed into PubMedBERT. The PubMedBERT embedding output was then used by the classifiers for predicting probabilities of the topic labels.
- The **BERTATT** model applies an attention of the PubMedBERT embedding output attending to the topic labels prior to the classifiers. The title and abstract of each article were concatenated and fed into PubMedBERT. The attention representations were computed, aggregated, and fed into the classifiers for topic label probability prediction.
- The **BERTMIL** model uses multi-instance learning to address the issue of large variation in the lengths of the articles, especially for articles with length exceeding the length limit of BERT models. In this model, abstract of each article was split into sentences. Title and the sentences were fed into PubMedBERT to output sentence embeddings which were then aggregated and used for topic label probability prediction.
- The **BERTBASEFOCAL** model replaces the Cross Entropy loss function used in the BERTBASE model with the Focal Loss (8) function to address the issue of imbalance in distribution of different topics.
- The **ENSEMBLE** model takes the topic label probabilities predicted by different models as input and computes an average of the predicted probabilities of each topic label as the final prediction.

### C. Experiment Settings

To train and validate the models, we split the training dataset into train and validation datasets at the ratio of 8:2. During the model development process, all models were trained with the train dataset, validated with the validation dataset, and evaluated with the development dataset. The models with best performance on the development dataset were used for prediction of the test dataset.

We fine-tuned the hyperparameters on the BERTBASE model and used the set of hyperparameters with best performance for all the models. We set the maximal sequence length at 512 for the models of BERTBASE, BERTATT, BERTBASEFOCAL, and 128 for the BERTMIL model. Each model was trained for maximal 20 epochs with learning rate of 1e-6 and batch size of 4. AdamW was used as the optimizer. The pooled output of the last layer of PubMedBERT was taken as the embedding for each article encoded by the model.

Both label-based and instance-based metrics of precision, recall and F1 score were used for evaluation of model performance. Evaluation results were calculated using the evaluation script provided. We used a deep learning model with TF-IDF as input as the baseline for evaluation of the development dataset. The baseline used for evaluation of the test dataset is the ML-Net (6), an end-to-end deep learning framework that combines the label prediction and label decision in the same network for multi-label biomedical text classification tasks.

## III. Results

Evaluation results of the development dataset and the test dataset are listed in Table III and Table IV respectively. The ensemble approach consistently produced the highest precision while the BERTBASEFOCAL model produced the highest recall in the label-based and instance-based metrics for both datasets. For F1 score, the ENSEMBLE approach achieved the best label-based micro average score, while the BERTBASEFOCAL model achieved the best label-based macro average score and the best instance-based score for the development dataset. For the test dataset, the ENSEMBLE approach consistently achieved the best F1 score in both label-based and instance-based metrics.

**TABLE III.** Evaluation results of the development dataset

| Models | P | R | F1 |
| --- | --- | --- | --- |
| **Label-based micro avg** |  |  |  |
| BERTBASE | 0.9119 | 0.8991 | 0.9055 |
| BERTATT | 0.9092 | 0.8974 | 0.9033 |
| BERTMIL | 0.9026 | 0.8921 | 0.8973 |
| BERTBASEFOCAL | 0.9086 | **0.9058** | 0.9072 |
| ENSEMBLE | **0.9183** | 0.8982 | **0.9081** |
| TF-IDF | 0.9029 | 0.8421 | 0.8715 |
| **Label-based macro avg** |  |  |  |
| BERTBASE | 0.8703 | 0.8488 | 0.8590 |
| BERTATT | 0.8651 | 0.8395 | 0.8515 |
| BERTMIL | 0.8630 | 0.8333 | 0.8470 |
| BERTBASEFOCAL | 0.8686 | **0.8563** | **0.8620** |
| ENSEMBLE | **0.8805** | 0.8408 | 0.8592 |
| TF-IDF | 0.8818 | 0.7371 | 0.7962 |
| **Instance-based** |  |  |  |
| BERTBASE | 0.9299 | 0.9253 | 0.9276 |
| BERTATT | 0.9281 | 0.9237 | 0.9259 |
| BERTMIL | 0.9184 | 0.9182 | 0.9183 |
| BERTBASEFOCAL | 0.9288 | **0.9308** | **0.9298** |
| ENSEMBLE | **0.9330** | 0.9250 | 0.9290 |
| TF-IDF | 0.8901 | 0.8724 | 0.8812 |

**TABLE IV.** Evaluation results of the test dataset

| Models | P | R | F1 |
| --- | --- | --- | --- |
| **Label-based micro avg** |  |  |  |
| BERTBASE | 0.9251 | 0.8814 | 0.9027 |
| BERTATT | 0.9284 | 0.8861 | 0.9067 |
| BERTMIL | 0.9141 | 0.8803 | 0.8969 |
| BERTBASEFOCAL | 0.9238 | **0.8880** | 0.9055 |
| ENSEMBLE | **0.9334** | 0.8841 | **0.9081** |
| Baseline (ML-Net) | 0.8756 | 0.8142 | 0.8437 |
| **Label-based macro avg** |  |  |  |
| BERTBASE | 0.8979 | 0.8328 | 0.8577 |
| BERTATT | 0.9062 | 0.8330 | 0.8636 |
| BERTMIL | 0.8917 | 0.8347 | 0.8518 |
| BERTBASEFOCAL | 0.9035 | **0.8394** | 0.8635 |
| ENSEMBLE | **0.9204** | 0.8345 | **0.8670** |
| Baseline (ML-Net) | 0.8364 | 0.7309 | 0.7655 |
| **Instance-based** |  |  |  |
| BERTBASE | 0.9358 | 0.9095 | 0.9225 |
| BERTATT | 0.9356 | 0.9131 | 0.9242 |
| BERTMIL | 0.9218 | 0.9062 | 0.9139 |
| BERTBASEFOCAL | 0.9337 | **0.9143** | 0.9239 |
| ENSEMBLE | **0.9380** | 0.9117 | **0.9247** |
| Baseline (ML-Net) | 0.8849 | 0.8514 | 0.8678 |

When examining the performance for each topic class, our evaluation results of the development dataset showed that more frequent topic classes, such as Prevention and Treatment, had higher scores, while less frequent topic classes, such as Transmission and Epidemic Forecasting, had lower scores.

## IV. Discussion

In addition to the models and strategies used to produce the submissions, we examined other possible strategies for tackling the challenges of the task. To address the issue of imbalance in distribution of each topic, we evaluated the feasibility of using data augmentation techniques, such as shuffling the order of sentences in abstract and randomly deleting a certain number of words in the article to increase the cases for less frequent topics. We observed that using data augmentation techniques was able to improve the precision score at the cost of decreasing recall for the corresponding topic classes. Further investigation might be worthwhile to achieve a better balance for the strategy.

Although our evaluation results showed that multi-instance learning (BERTMIL model) was not able to achieve better performance, training different models for articles with different lengths may have the potential to improve the performance since there is a large variation in the article lengths in the dataset.

## V. Conclusion

During our participation of BioCreative VII track 5, we tackled the multi-label topic classification for automated annotation of COVID-19 literature by evaluating several deep learning models of different architectures. Evaluation on the development and test datasets showed consistent results that the ENSEMBLE approach achieved the highest F1 score and precision and the BERTBASEFOCAL model achieved the highest recall in both label-based and instance-based metrics. Compared to the performance of the baseline model, our models were able to achieve significantly better performance. The performance of our models also compared favorably to those of other participating teams.

